# GraphBG: Fast Bayesian Domain Detection via Spectral Graph Convolutions for Multi-slice and Multi-modal Spatial Transcriptomics

**DOI:** 10.64898/2026.03.28.715026

**Authors:** Van Hoan Do, Thi Phuong Lan Tran, Stefan Canzar

## Abstract

Spatial transcriptomics (ST) technologies enable measurement of gene expression with spatial context, offering unprecedented insight into tissue architecture and cellular microenvironments. A fundamental analysis task is the identification of spatial domains, i.e., contiguous regions with distinct molecular profiles. As ST datasets scale to larger tissue areas, multiple slices, and multiple molecular modalities, there is a growing need for clustering methods that are accurate, scalable, and capable of integrating diverse spatial and molecular signals. We present *GraphBG*, a unified and scalable framework for spatial domain detection in ST data. GraphBG integrates approximate spectral graph convolutions with a variational Bayesian Gaussian mixture model, enabling robust representation learning and clustering of spatially coherent domains. We extend this core model to support multi-slice analysis (*GraphBG-MS*) through metacell aggregation, batch correction, and joint clustering, and to multi-modal spatial omics data (*GraphBG-MM*) via modality-specific graph encodings and kernel canonical correlation analysis. Across diverse real and simulated datasets, GraphBG consistently outperforms existing methods in domain coherence, scalability, and biological interpretability. Notably, it accurately clusters over 370,000 cells from 31 MERFISH tissue slices in just 5 minutes and integrates spatial transcriptomic and proteomic data for improved domain resolution. Applying GraphBG-MS to mouse liver ST data, we show that it captures canonical lobular zonation and disease-specific remodeling, highlighting its ability to reveal biologically meaningful tissue organization.

## 1. Introduction

Spatial transcriptomics (ST) technologies have revolutionized our ability to study gene expression within its spatial context, providing new insights into tissue organization, development, and disease [1–3]. Unlike traditional single-cell RNA sequencing, ST preserves the spatial coordinates of gene expression measurements, enabling the exploration of spatially regulated biological processes and cell-cell interactions [4, 5]. This capacity has proven indispensable for constructing high-resolution tissue atlases, uncovering structural compartments, and identifying spatial gradients of transcriptional activity. As these technologies continue to scale, the ability to analyze ST data at large scale, across hundreds of thousands of spots, multiple tissue slices, or entire organs, has become essential for realizing their full potential [6–8].

A fundamental task in spatial transcriptomics analysis is the identification of *spatial domains*, i.e.groups of spatially contiguous regions with coherent transcriptional profiles [9–11]. Spatial domain detection enables reconstruction of anatomical structures, inference of functional regions, and discovery of domain-specific marker genes. It also facilitates downstream tasks such as trajectory inference [12], spatially resolved gene regulation [13], and disease subtype classification. Accurate and scalable spatial clustering methods are therefore central to the ST analysis pipeline.

Early attempts at spatial clustering relied on non-spatial methods such as Louvain [14] and Seurat [15], which considered only gene expression and often produced fragmented or biologically implausible domains. More recent approaches incorporate spatial information to improve domain coherence. SpaGCN [16], stLearn [12], and BayesSpace [17] use graph-based or probabilistic models to integrate spatial proximity with gene expression. STAGATE [13] and CCST [18] employ graph neural networks to learn spatial embeddings. SpaceFlow [19] and GraphST [20] utilize self-supervised learning strategies to encode spatial context without requiring annotations. For multi-slice integration, methods such as SpaceFlow-DC [9] apply the SpaceFlow model independently to each slice and align embeddings post hoc using batch correction tools like Harmony [21].

Despite recent progress, current spatial clustering methods face several limitations. First, scalability remains a major challenge. Early spatial clustering tools were developed for datasets with fewer than 10,000 cells or spots, reflecting the capabilities of earlier technologies. However, modern platforms such as Slide-seqV2 now produce datasets with tens or even hundreds of thousands of spots [9], exposing severe memory and runtime bottlenecks in many existing methods. Second, most tools are designed for single-slice analysis and cannot effectively integrate information across multiple tissue sections [9, 10]. This limitation becomes problematic as an increasing number of studies generate multi-slice spatial transcriptomics data to construct large-scale spatial atlases. Independent clustering of each slice often results in inconsistent or incomparable domain labels and clustering granularities, especially in continuous tissues with subtle boundaries. These inconsistencies hinder data integration and downstream analyses, such as identifying condition- or patient-specific spatial features [9, 10]. Finally, current methods generally assume unimodal input and lack support for emerging multi-omics spatial datasets that jointly measure gene expression, protein abundance, or chromatin accessibility [22]. As multi-modal spatial profiling becomes more common, there is an urgent need for integrative methods that can leverage complementary information across modalities for more accurate and robust spatial domain detection.

To address these limitations, we introduce *GraphBG*, a unified and scalable framework for spatial domain detection in spatial transcriptomics (ST) data. GraphBG combines spatially informed spectral graph convolutional encoding with a variational Bayesian Gaussian mixture model (VB-GMM) to enable accurate, interpretable, and scalable clustering. The use of approximate spectral graph convolutions efficiently captures local spatial dependencies, while the variational Bayesian formulation provides robust, uncertainty-aware clustering and mitigates overfitting. Together, these components allow GraphBG to overcome both computational and modeling challenges in modern ST analysis. To support large-scale analysis across multiple tissue sections, GraphBG performs slice-wise preprocessing and metacell aggregation to reduce complexity, followed by batch correction with ComBat and joint clustering via VB-GMM. For multi-modal ST integration, GraphBG learns modality-specific graph embeddings, which are aligned into a shared latent space using kernel canonical correlation analysis (KCCA). Clustering in this unified space using VB-GMM enables accurate domain detection using signals from each molecular modality.

We benchmarked our methods against leading spatial clustering approaches across diverse real and simulated datasets, including large-scale and multi-slice collections. GraphBG supports unimodal, multi-slice, and multi-modal analysis, enabling accurate spatial domain detection, cross-slice alignment, and integration of transcriptomic and proteomic modalities. Together, these capabilities provide a unified and scalable framework for spatial clustering across a wide range of experimental settings.

## 2. Results

### 2.1. Overview of our methods

We introduce GraphBG, a unified and scalable framework for spatial domain detection in spatial transcriptomics (ST) data, supporting unimodal, multi-modal, and multi-slice analyses. GraphBG integrates graph-based representation learning with Bayesian statistics for clustering, offering a principled approach to capture both gene expression variability and spatial structure.

For unimodal ST data, GraphBG begins with standard normalization and preprocessing, followed by the application of approximate spectral graph convolutions to encode spatial dependencies among spots or cells. The learned spatially-informed embeddings are then clustered using a variational Bayesian Gaussian mixture model (VB-GMM) [23]. A final refinement step improves spatial coherence by smoothing cluster assignments based on local spatial neighborhoods. The pipeline is given in Fig. 1 (top).

**Figure 1.**
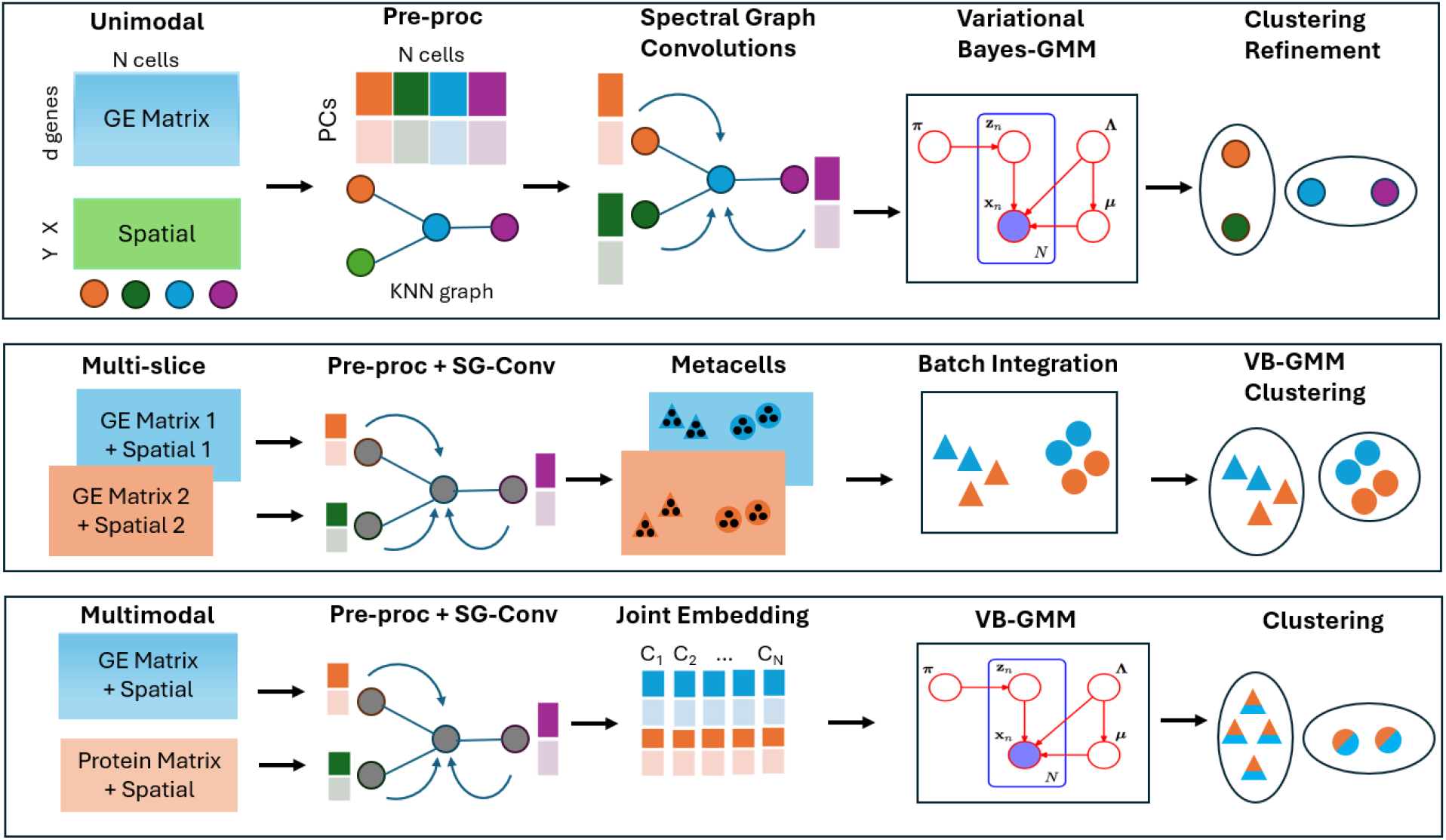
The overall backbone framework of GraphBG. The main steps were included in carrying out spatial domain detection from the input spatial omics data: *top:* Unimodal ST data, *middle:* multi-slice ST data (colors indicate slices, shapes indicate spatial domains), *bottom:* multi-modal ST data (colors indicate modalities, shapes indicate spatial domains).

To handle large-scale datasets comprising multiple tissue slices, we propose GraphBG-MS, a strategy that computes metacells per slice using minibatch k-means clustering. The metacell embeddings, derived from averaged graph representations, are batch-corrected using ComBat [24], and then jointly clustered via VB-GMM. Metacell-level cluster labels are then propagated back to individual cells or spots (Fig. 1 (middle)).

For multi-modal ST data, we introduce GraphBG-MM, which incorporates multi-view graph representations across modalities (e.g., gene expression and protein abundance). Modality-specific embeddings are aligned using kernel canonical correlation analysis [25, 26] before VB-GMM clustering. This enables integration of complementary signals while leveraging spatial information (Fig. 1 (bottom)).

### 2.2. GraphBG’s accurately detects spatial domains in single slice

To ensure consistency and comparability with prior benchmarking studies, we adhered to the evaluation methods and metric definitions established by Yuan et al. [9]. To compare the spatial clustering performance of GraphBG with existing methods, we selected six representative tools from three methodological categories: (1) a traditional single-cell clustering method, Louvain [14]; (2) Bayesian approaches including BASS [27] and BayesSpace [17]; and (3) graph neural network (GNN)-based deep learning methods GraphST [20], SCAN-IT [28], and SpaceFlow [19]. These methods were selected because they rank among the top performers in terms of both accuracy and computational efficiency in recent benchmarks [9]. Consistent with the benchmark [9], we employed three complementary clustering evaluation metrics: Normalized Mutual Information (NMI), which measures the overall agreement between predicted and reference labels; Homogeneity (HOM), which favors methods that assign each cluster to a single ground-truth class; and Completeness (COM), which favors methods that group together all samples of the same class. These metrics are particularly appropriate for spatial clustering as they jointly capture clustering accuracy, purity, and coverage of annotated tissue domains. Our evaluation was conducted on the gold-standard 10x Visium human dorsolateral prefrontal cortex (DLPFC) dataset [29], which comprises spatial transcriptomic profiles from 12 annotated tissue slices, each containing measurements for over 20,000 genes and a mean of 3,844 spots.

As shown in Fig. 2, GraphBG consistently performed well across evaluation metrics. In particular, it reached the highest mean NMI (0.692) and HOM (0.711), while maintaining a competitive COM (0.676). Compared with GraphST, which achieved the highest COM (0.692), GraphBG reached comparable NMI (0.692 vs. 0.673) and moderately higher HOM (0.711 vs. 0.657). Among Bayesian approaches, BayesSpace obtained intermediate values across all three metrics (NMI 0.601, HOM 0.614, COM 0.590). SpaceFlow achieved moderate HOM (0.656), comparable to GraphST but lower than GraphBG, while its NMI (0.496) and COM (0.402) were substantially weaker. The Louvain algorithm, which does not incorporate spatial information, consistently underperformed (mean scores only slightly above 0.20 across all metrics), emphasizing the value of spatial context in tissue segmentation. Overall, GraphBG demonstrates a moderate yet consistent improvement over strong GNN-based baselines across NMI, HOM, and COM, while also outperforming Bayesian and traditional clustering methods.

**Figure 2.**
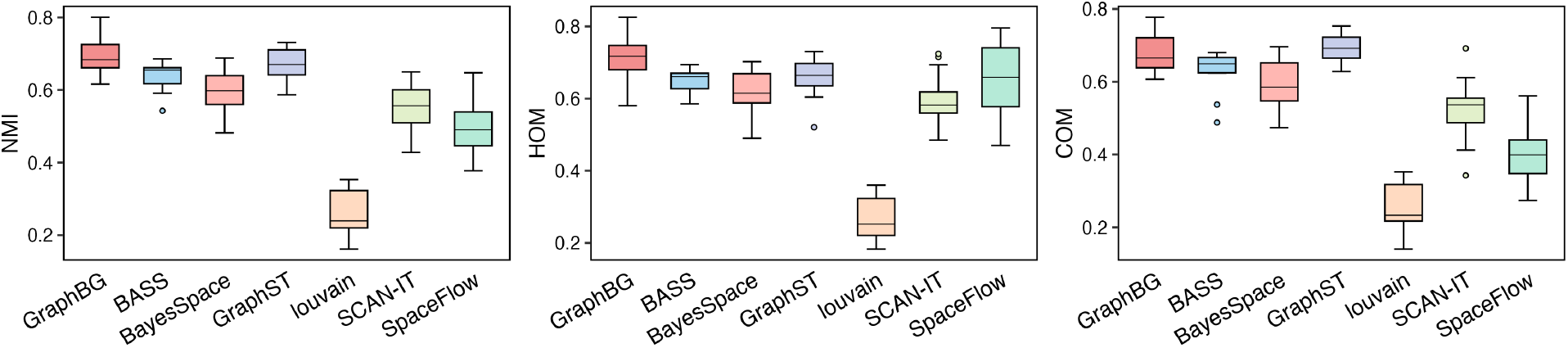
Performance of GraphBG and competitive methods on the 10x Visium dataset.

### 2.3. Benchmark analysis on various technologies

After establishing the strong performance of GraphBG on the 10x Visium dataset, we next evaluated its generalizability to various sequencing platforms: 10x Visium, BaristaSeq [30], MERFISH [31], osmFISH [32], STARmap [33], and a 1,000-gene subset of STARmap (STARmap*) [33]. These datasets offer manually curated domain annotations suitable for benchmarking. GraphBG was benchmarked against the same set of spatial clustering methods as in previous experiments, excluding BayesSpace due to its incompatibility with imaging-based datasets. Across all datasets and evaluation metrics (NMI, HOM, COM), GraphBG consistently achieved the highest mean scores (NMI ∼0.65, HOM ∼0.69, COM ∼0.61) (Fig. 3). Among GNN-based methods, GraphST was the closest competitor, followed by SCAN-IT and SpaceFlow whose accuracy was consistently lower than GraphBG’s. SpaceFlow showed larger fluctuations particularly in terms of completeness. Bayesian models such as BASS also scored lower than GraphBG in NMI and homogeneity, while the non-spatial Louvain algorithm performed markedly worse. Scores only slightly above 0.20 underscore the necessity of incorporating spatial information. Collectively, these findings demonstrate that GraphBG achieves consistently high and reliable clustering accuracy, with small yet robust improvements over current baseline approaches.

**Figure 3.**
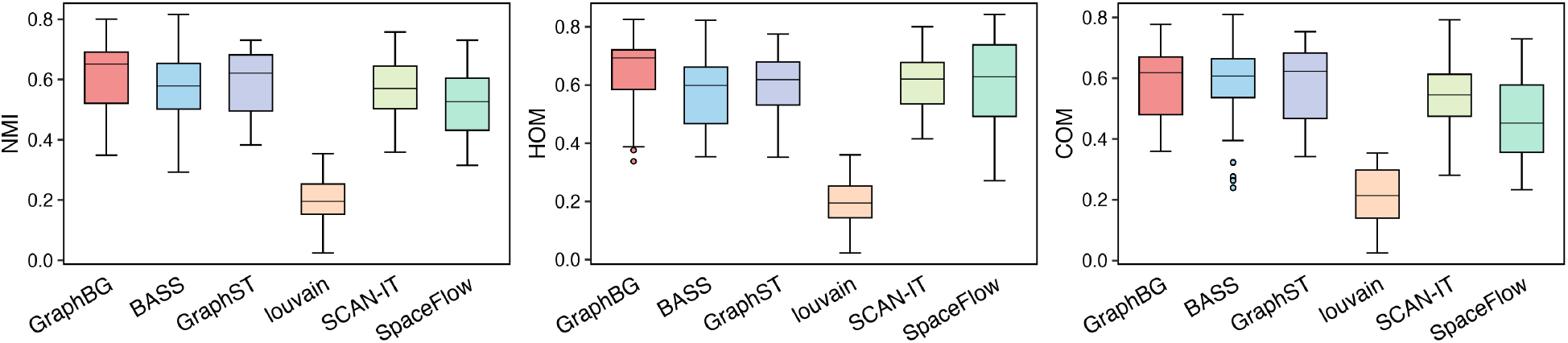
Performance of GraphBG and competitive methods on six spatial transcriptomics datasets.

### 2.4. GraphBG-MM utilizes multiple modalities more effectively

We demonstrated the capability of GraphBG-MM to leverage complementary information from multi-modal data to more accurately refine spatial domains in spatial multi-omics datasets. Specifically, we re-analyzed both publicly available and simulated datasets generated using scDesign3 [34]. We compared our method to SpatialGlue [22], a graph neural network based model for domain detection from spatial multi-omics. We created six multi-modal spatial transcriptomics datasets by simulating CITE-seq profiles [35] with scDesign3 and assigning them to brain regions defined by an anatomical atlas [36] (Supplementary Figure 1). scDesign3 simulated RNA and ADT modalities based on a reference dataset with known cell types [35]. Spatial coordinates were sampled from annotated brain regions, with cell types assigned according to region-specific probabilities. The resulting datasets preserved the multi-modal structure of CITE-seq and provided ground-truth spatial domain labels for benchmarking. We systematically varied two key parameters—the number of cells (n = 2000, 5000, 10000) and the number of measured genes (p = 100 or 1000)—to mimic targeted versus untargeted assays.

We first investigated if integrating RNA and protein is crucial for accurate spatial domain detection. To this end, we evaluated GraphBG-MM under three input configurations: using only RNA, using only protein, and using both modalities together (RNA + protein). As shown in Supplementary Figure 2, the multi-modal configuration consistently outperformed both single-modality configurations across all three evaluation metrics. The gains were particularly evident in larger datasets, where RNA + protein (multi-modal) achieved HOM, COM, and NMI values close to 0.7, while RNA-only and protein-only configurations remained around and 0.55, respectively. These results confirm that RNA and protein carry complementary information, and their integration enables GraphBG-MM to more accurately capture spatial domain structures.

After establishing the advantage of multi-modal integration, we compared GraphBG-MM against SpatialGlue, a graph neural network-based model for spatial multi-omics. Across all simulation conditions, GraphBG-MM consistently outperformed SpatialGlue in all three metrics, as visible in the aggregated trend (Fig. 4a) and further detailed in the dataset-level results (Supplementary Figure 3). Importantly, the performance gap became more pronounced in scenarios with increased cell count and gene number. When the number of cells rose from 2000 to 10000, we observed substantial gains in HOM, COM, and NMI for both methods; however, GraphBG-MM maintained a clear advantage. For instance, in the largest dataset (n=10000, p=1000), GraphBG-MM achieved HOM, COM, and NMI values of around 0.7, while SpatialGlue plateaued at approximately 0.6. Similarly, as the number of genes increased from 100 to 1000, both methods showed improved clustering performance, with GraphBG-MM benefiting more substantially, especially in terms of NMI.

**Figure 4.**
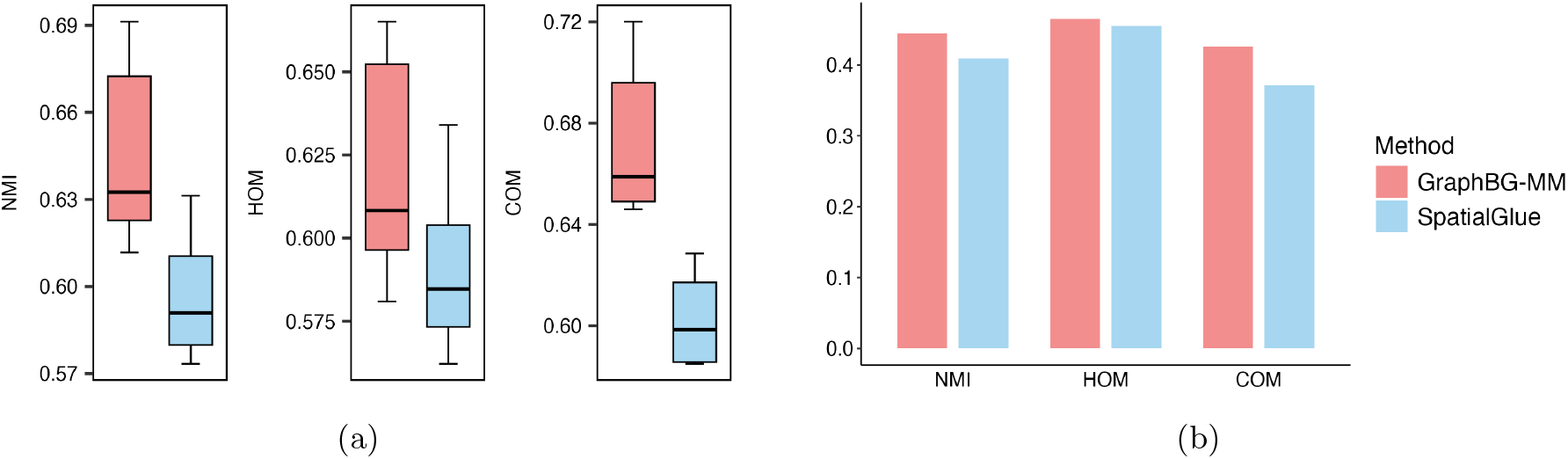
NMI, homogeneity, and completeness of domains detected by GraphBG-MM and SpatialGlue on six simulated datasets (a) and in a human lymph node dataset (b).

We next benchmarked GraphBG-MM and SpatialGlue on a real human lymph node dataset generated with 10x Genomics Visium spatial transcriptomics and proteomics co-profiling, using hematoxylin and eosin (H&E) annotations from the original study as ground truth [22]. On this dataset, GraphBG-MM again demonstrated good performance across all supervised metrics, improving NMI (0.45 vs 0.41) and completeness (0.43 vs 0.37) compared to SpatialGlue (Fig. 4b).

Next, we assessed the spatial coherence captured by GraphBG-MM by benchmarking it against SpatialGlue on four thymus spatial profiling datasets (mouse_thymus_1–4 in Fig. 5). These datasets were generated using two distinct platforms: Stereo-CITE-seq [37] and SPOTS [38]. In the absent of manual annotation, we followed [9] and employed Moran’s I to quantify spatial coherence, a widely used metric that measures global spatial autocorrelation by assessing whether similar values (in our case, cluster labels) tend to occur near each other in space. A higher Moran’s I score indicates stronger spatial autocorrelation and thus better spatial consistency of clustering results. Our implementation of Moran’s I follows the procedure and code released in the CORAL study [39]. Across all datasets, GraphBG-MM achieved markedly higher Moran’s I scores compared to SpatialGlue, highlighting its superior ability to preserve spatial autocorrelation (Fig. 5). Notably, in the fourth dataset mouse_thymus_4 (Fig. 5), GraphBG-MM maintained a high Moran’s I score of 0.745, whereas SpatialGlue dropped to just 0.161.

**Figure 5.**
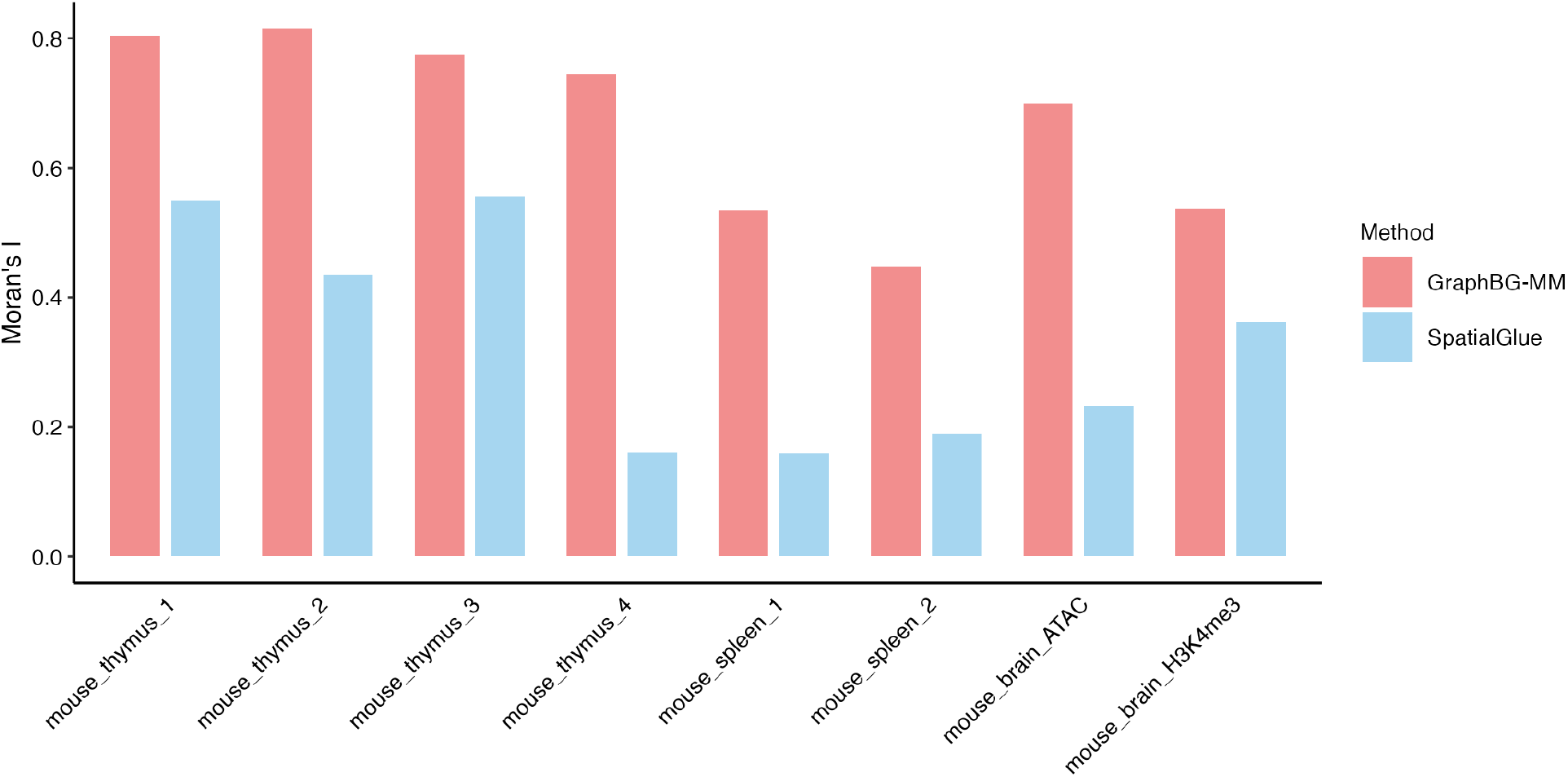
Comparison of Moran’s I scores for spatial clustering by GraphBG-MM and SpatialGlue across four thymus datasets and two spleen datasets.

We next applied GraphBG-MM to two mouse spleen spatial multi-omic datasets (mouse_spleen_1 and mouse_spleen_2 in Fig. 5), consisting of whole transcriptomes and surface protein profiles acquired using the SPOTS protocol with Visium technology. GraphBG-MM consistently achieved substantially higher Moran’s I scores (0.538 vs 0.157 for mouse_spleen_1; 0.448 vs 0.181 for mouse_spleen_2) (Fig.5), indicating stronger spatial autocorrelation compared to SpatialGlue.

In a final evaluation, we assessed GraphBG-MM on two spatial multi-omic datasets from postnatal day (P)22 mouse brain coronal sections (Fig.5). The first dataset (mouse brain ATAC) simultaneously captures chromatin accessibility and gene expression using spatial ATAC–RNA-seq, while the second dataset (mouse brain H3K4me3) profiles transcriptomes alongside H3K4me3 histone modifications via CUT&Tag. As shown in Fig. 5, GraphBG-MM consistently outperformed SpatialGlue across both brain datasets. In the mouse_brain_ATAC dataset, GraphBG-MM achieved a Moran’s I score of 0.699, substantially higher than the 0.232 score obtained by SpatialGlue. Similarly, in the mouse_brain_H3K4me3 dataset, GraphBG-MM reached a score of 0.537, compared to 0.362 for SpatialGlue.

### 2.5. GraphBG-MS: multi-slice spatial clustering analysis on a large-scale dataset

To address the challenge of identifying coherent spatial domains across multiple tissue slices, we introduce *GraphBG-MS*. The method is designed to scale spatial clustering to large multi-slice datasets while maintaining alignment across slices and preserving biological structure (see Methods section).

We compared our method to *SpaceFlow-DC* [9], which adopts a divide-and-conquer strategy to speed up spatial clustering across multiple slices. In SpaceFlow-DC, the original SpaceFlow model is applied independently to each tissue slice to generate slice-specific embeddings. These embeddings are then integrated using Harmony [21], a batch correction algorithm that aligns shared structure across slices. Final clustering is performed on the harmonized embedding using graph-based algorithms such as Louvain, Leiden, or Gaussian mixture models (e.g., `mclust`). Compared to SpaceFlow-DC, GraphBG-MS incorporates an explicit probabilistic model in the form of VB-GMM for joint clustering. Furthermore, it uses metacells to improve scalability and robustness. We also compared against *scNiche* [40], a recently developed method that models cell niches by integrating multi-views features of the cell, including both intrinsic molecular profiles and microenvironmental features such as neighborhood composition.

We conducted a comparison between GraphBG-MS, SpaceFlow-DC, and scNiche using a large-scale multi-slice MERFISH dataset comprising 31 tissue slices and over 300,000 cells [41]. Both SpaceFlow-DC and scNiche are designed to overcome the challenges of spatial clustering in multi-slice settings by leveraging inter-slice or multi-view information. However, our results show that GraphBG-MS significantly outperforms both competitors across multiple evaluation metrics. Specifically, GraphBG-MS achieved a NMI score of 0.71, a HOM score of 0.65, and a COM score of 0.77, compared to 0.59 (NMI), 0.57 (HOM), and 0.61 (COM) for SpaceFlow-DC, and 0.66 (NMI), 0.64 (HOM), and 0.69 (COM) for scNiche (Fig. 6). In terms of runtime, GraphBG-MS completed in approximately 5 minutes, significantly faster than both SpaceFlow-DC (133 minutes) and scNiche (221 minutes) under the same hardware conditions.

**Figure 6.**
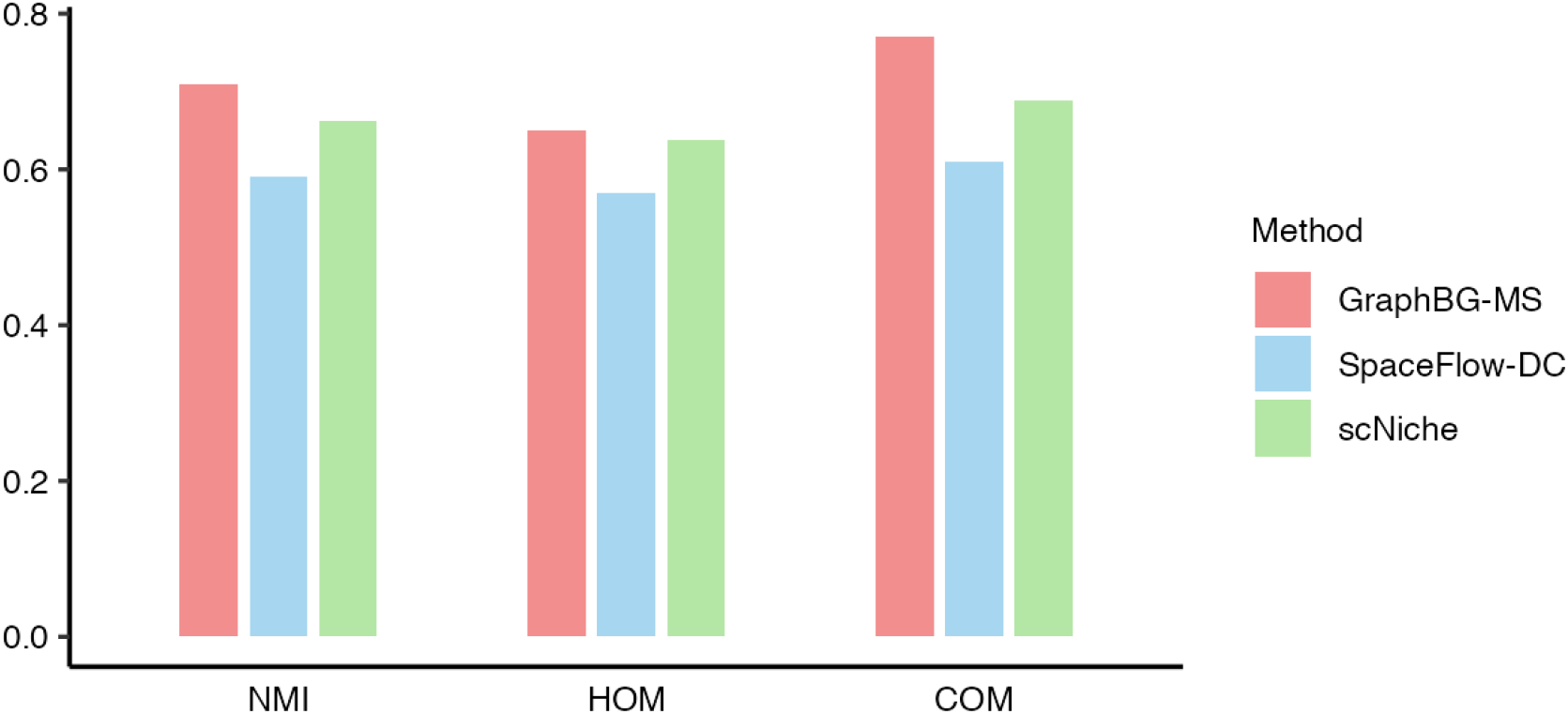
Comparison of spatial clustering performance among GraphBG-MS, SpaceFlow-DC, and scNiche on a large-scale multi-slice MERFISH dataset (31 slices, *>*300,000 cells).

Together, these results suggest that GraphBG-MS provides more consistent and accurate clustering that aligns better with expert annotations, while maintaining substantial computational efficiency.

### 2.6. GraphBG-MS captures liver zonation and disease-specific spatial domain remodeling

To demonstrate the biological relevance of domains detected across multiple slices of normal and diseased tissue, we re-analyzed high-resolution spatial transcriptomic profiles from mouse livers generated with Seq-Scope [42]. The dataset comprised six normal samples and four cases of early-onset liver failure driven by excessive mTORC1 signaling (TD models), and had previously been examined with scNiche [40]. GraphBG-MS identified 18 spatial domains (niches) (Supplementary Figure 4) and we compared them directly to the domains reported by scNiche (Supplementary Figure 5).

To validate the performance of GraphBG-MS, we adopted a strategy similar to that of [40], examining zonation patterns in normal livers along the central-to-portal axis (Domains 8, 9, 14, and 12) (Fig. 7-a). Differential expression analysis across these domains revealed clear spatial trends that aligned with the established lobular architecture. For example, from Domain 8 to Domain 12, the expression of pericentral genes (*Cyp2e1, Cyp1a2, Mup17, Gsta3, Gulo*) gradually decreased, whereas the expression of periportal genes (*Ass1, Alb, Cyp2f2, Sds, Mup20*) showed a progressive increase [40, 43–47] (Supplementary Figures 6). To benchmark the ability of GraphBG-MS against scNiche in recovering liver zonation programs, we compared the gene signatures distinguishing Domains 8 and 9 (encompassing major pericentral hepatocytes) from Domains 14 and 12 (encompassing major periportal hepatocytes) (Supplementary Figure 7, Supplementary Table 1, Fig. 7-b). The two methods shared a large core set of canonical zonation genes, including urea cycle and amino acid metabolism markers (*Cps1, Ass1*), classic cytochrome P450s (*Cyp2e1, Cyp2c29/37/50/67/70*), and oxidative stress enzymes (*Gpx3, Gstp1, Fmo5*) [43–47]. However, the two approaches differed substantially in their coverage of canonical metabolic pathways. GraphBG-MS outperformed scNiche, as it recovered a broad set of cytochrome P450 enzymes (*Cyp17a1, Cyp2b9, Cyp3a44*) (Supplementary Table 1, Fig. 7-c), thereby highlighting the classical role of the pericentral zone in xenobiotic metabolism. In contrast, scNiche detected only a narrower spectrum of P450 genes, with a stronger bias toward Mup family members. This demonstrates that GraphBG-MS provides a more comprehensive and balanced representation of pericentral functions by capturing both cytochrome P450 enzymes and Mup family members. Similarly, for periportal hepatocytes, GraphBG-MS successfully recovered hallmark metabolic genes involved in retinol metabolism (*Fabp1, Rdh7*) and the urea cycle (*Arg1*) (Supplementary Table 1, Fig. 7-c). By comparison, scNiche primarily highlighted enrichment for secreted plasma proteins (*Cp, F2, Fn1*), reflecting a narrower view of periportal hepatocytes’ role in systemic protein synthesis.

**Figure 7.**
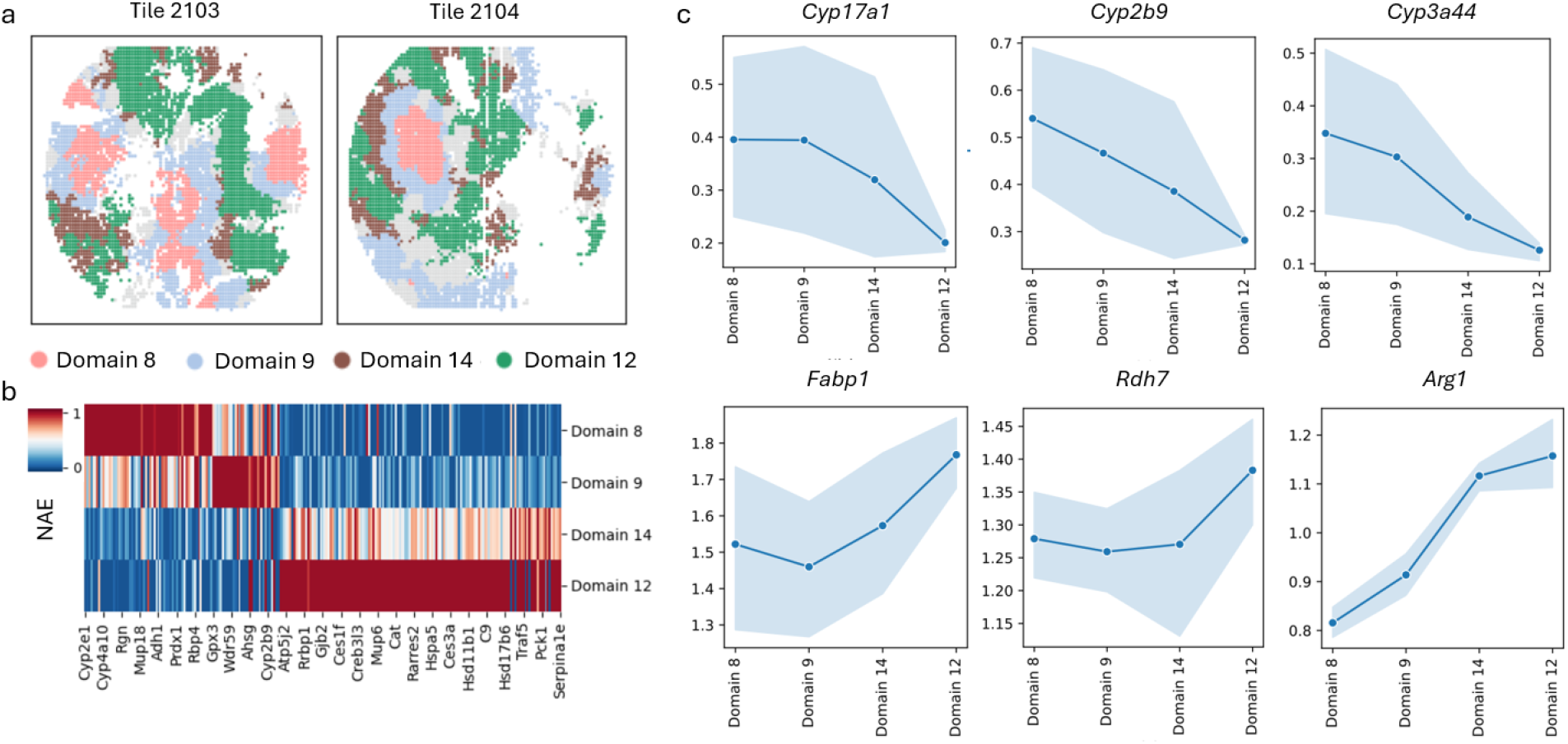
GraphBG-MS reveals the molecular architecture of normal liver zonation. **a** Domains identified by GraphBG-MS in tiles 2103 and 2104. Cells are colored by the domain labels. **b** Normalized average expression (NAE) of 235 genes differentially expressed (adjusted p-value *<* 0.05) across Domains 8, 9, 14, 12. The complete gene list is given in Supplemental Table 1. Statistical significance was assessed using a two-sided Wilcoxon rank-sum test, with multiple testing controlled by the Benjamini-Hochberg procedure. **C** Average expression levels of *Cyp17 a1, Cyp2b9, Cyp3a44, Fabp1, Rdh7*, and *Arg1* across Domains 8, 9, 14, 12. Plots show mean expression levels, with error bars indicating 95% confidence intervals.

Finally, we compared the spatial domains in the portal node between normal livers (Domain 12) and TD livers (Domain 10) (Fig. 8-a). Differential expression analysis revealed injury-associated transcriptional changes, largely overlapping with those observed in scNiche (Fig. 8-b, Supplementary Figure 8). Specifically, scNiche highlighted hepatocyte-centric stress responses, including reactive oxygen species detoxification (*Gpx3*) and altered lipogenesis (*Scd2*), together with cytochrome P450 enzymes (Supplementary Figure 8). These signatures align with direct mTORC1-driven metabolic stress in hepatocytes. In contrast, GraphBG-MS emphasized systemic and fibrotic responses, capturing glutathione detoxification (*Gsta1*), acute-phase signaling (*Orm3*), and extracellular matrix remodeling (*Col5a2*) (Fig. 8-b,c). This reflects downstream progression from primary hepatocyte stress toward inflammation and fibrosis. Thus, scNiche provides a lens on early metabolic rewiring and oxidative stress, whereas GraphBG-MS reveals the broader systemic and fibrotic consequences of mTORC1 hyperactivation.

**Figure 8.**
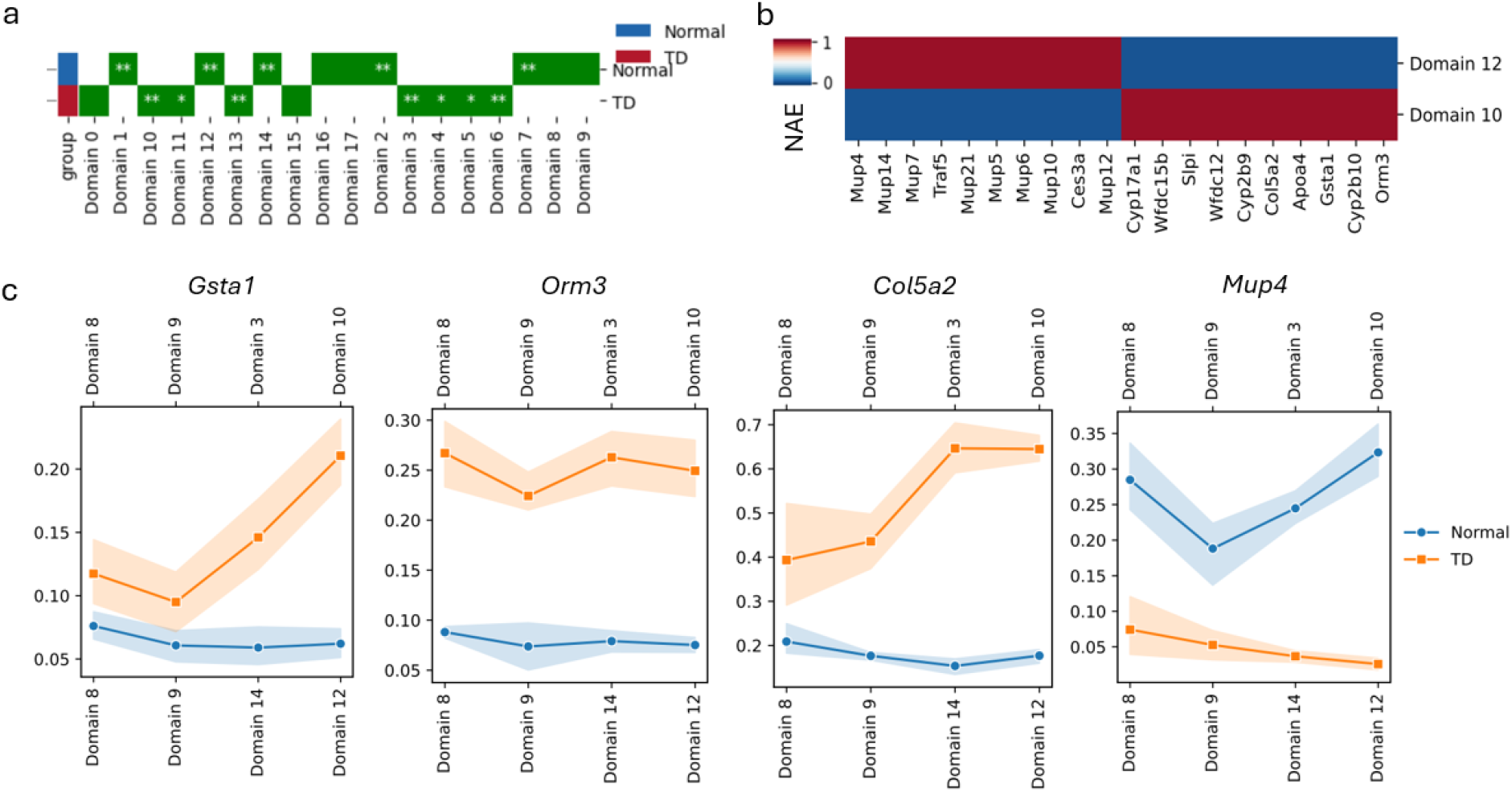
GraphBG-MS identifies liver injury spatial domains (niches) and tracks remodeling from normal to failure. **a** Domain-wise enrichment scores comparing normal and TD donors. Statistical significance was assessed using a one-sided Mann–Whitney U test. (∗ *p*≤ 0.05; ∗∗*p* ≤0.01). **b** Normalized average expression (NAE) of top 20 differentially expressed genes (adjusted p-value *<* 0.05) in Domain 10 and Domain 12. **C** Comparative analysis of gene expression gradients from central vein to portal node domains in normal versus TD livers. Plots show mean expression levels, with error bars indicating 95% confidence intervals.

### 2.7. Runtime comparison of GraphBG and competing methods in a single-slice analysis

We evaluated the computational efficiency of GraphBG against state-of-the-art methods GraphST and SpaceFlow in a single-slice spatial transcriptomics setting. Experiments were conducted on the MERFISH mouse brain from the Brain Aging Spatial Atlas [41], comprising approximately 378,000 cells. We concatenate all tissue slices into a unified dataset and append an additional coordinate or slice identifier to each spot to preserve slice-specific information. To assess scalability, we subsampled the dataset at different sizes (10k, 50k, and 100k cells), and also evaluated the full dataset when possible. All methods were executed under identical hardware conditions, and runtime was measured in minutes.

**Table 1:** Runtime comparison (in minutes) on varying dataset sizes. “–” indicates failure due to memory error.

| # Cells | GraphST | GraphBG | SpaceFlow |
| --- | --- | --- | --- |
| 10k | 4.06 | 0.22 | 2.21 |
| 50k | 75.93 | 2.99 | 17.11 |
| 100k | — | 8.33 | 23.82 |
| Full (378k) | — | 40.66 | 106.17 |

GraphBG demonstrates superior scalability, with significantly faster runtimes across all dataset sizes. At the 50k cell level, GraphBG completes in under 3 minutes, whereas GraphST requires over 75 minutes and SpaceFlow takes more than 17 minutes. Notably, GraphST fails to complete at 100k cells and beyond due to memory limitations (315 Gb), while GraphBG processed the full 378k-cell dataset in just 40.66 minutes.

Although SpaceFlow completes execution at all scales, it suffers from substantially lower accuracy in our previous benchmarking study. Specifically, on the gold-standard 10x Visium dataset, SpaceFlow’s average Normalized Mutual Information (NMI) is below 0.5, while GraphBG achieves an NMI of approximately 0.7. Furthermore, benchmark across different spatial transcriptomics technologies, the average NMI of SpaceFlow remains below 0.52, compared to GraphBG’s average NMI of around 0.62. These results underscore the trade-off in SpaceFlow between runtime and clustering accuracy, and emphasize GraphBG’s robustness in both performance and precision across diverse platforms.

## 3. Discussion

Spatial transcriptomics has transformed our understanding of tissue organization by enabling transcriptome-wide profiling with spatial resolution. A critical task in analyzing such data is the identification of spatial domains-discrete tissue regions with coherent gene expression and spatial structure-offering insights into tissue architecture, disease microenvironments, and cellular domains. Large-scale efforts are now underway to construct comprehensive spatial atlases across organs, species, and developmental stages. However, current spatial domain detection methods face substantial limitations. Most notably, scalability remains a bottleneck, particularly as datasets now routinely include hundreds of thousands of cells across dozens of slices. Furthermore, existing methods often fail to effectively integrate across batches or slices, and are not designed to handle multi-modal spatial omics data that jointly profile gene expression, chromatin states, or protein abundance. To address these challenges, we developed GraphBG, a general and scalable framework for spatial domain detection, and extended it to two complementary settings: GraphBG-MS for multi-slice integration, and GraphBG-MM for spatial multi-omics clustering.

Across diverse benchmarks, our method consistently outperforms existing tools in accuracy, spatial coherence, and computational efficiency. In the unimodal setting, GraphBG achieved the highest scores across key evaluation metrics (NMI = 0.68, HOM = 0.71) on the gold-standard 10x Visium DLPFC dataset, outperforming 13 leading methods including GraphST, BayesSpace, BASS, and SpaceFlow. When applied to six spatial transcriptomics technologies covering distinct biological systems, GraphBG maintained superior accuracy and robustness, confirming its generalizability. For spatial multi-omics analysis, GraphBG-MM demonstrated clear advantages over SpatialGlue in both simulation and real data, consistently achieving higher performance across all metrics. In particular, GraphBG-MM better preserved spatial autocorrelation as measured by Moran’s I, especially in large and complex tissues such as spleen and postnatal mouse brain. In the multi-slice setting, GraphBG-MS surpassed the divide-and-conquer strategy SpaceFlow-DC by a substantial margin (NMI: 0.71 vs. 0.59) while reducing runtime by over 25-fold on a 31-slice MERFISH dataset (*>*300,000 cells). Finally, application of GraphBG-MS to mouse liver data showed that it faithfully recapitulates canonical lobular zonation and uncovers disease-specific spatial remodeling under mTORC1-driven liver failure, underscoring the framework’s ability to capture biologically meaningful tissue organization across both normal and pathological states.

Several key design features of GraphBG contribute to its superior performance. First, the use of spatially informed graph construction ensures that both transcriptomic similarity and spatial proximity are leveraged effectively. Second, the usage of approximate spectral graph convolution and metacell-based abstraction reduces computational burden while enhancing robustness, making the framework highly scalable to large datasets. Third, the use of variational Bayesian Gaussian mixture modeling (VB-GMM) enables principled probabilistic clustering that accommodates uncertainty. Fourth, GraphBG-MS introduces cross-slice integration through shared embedding and joint modeling, while GraphBG-MM incorporates multi-modal fusion without sacrificing spatial coherence.

Despite its strengths, GraphBG opens up several avenues for future work. A natural extension is to incorporate temporal information or trajectory dynamics in spatiotemporal transcriptomics. Likewise, extending GraphBG-MM to integrate spatial imaging data (e.g., histopathology or spatial metabolomics) alongside molecular profiles could enhance spatial context understanding. From a modeling perspective, integrating hierarchical or multi-resolution clustering within the probabilistic framework may offer finer-grained annotations of complex tissues. Finally, building interactive, interpretable tools based on GraphBG could facilitate adoption by biologists and pathologists, and support applications in tissue atlasing, developmental biology, and clinical diagnostics.

In summary, we present GraphBG as a general, scalable, and extensible framework for spatial domain detection in spatial transcriptomics and spatial multi-omics. Through rigorous benchmarking, we demonstrate that GraphBG, along with its extensions GraphBG-MS and GraphBG-MM, consistently delivers good accuracy, spatial coherence, and scalability across a wide range of data types and biological systems. Our framework addresses major limitations in current spatial clustering methods and provides a powerful tool for the next generation of spatial omics analysis at scale.

## 4. Methods

### 4.1. GraphBG: Graph-based Bayesian Gaussian mixture model for spatial domain detection of unimodal ST data

We consider gene expression counts *X* ∈ ℝ^*N ×d*^, where *N* is the number of spots or cells, and *d* is the number of features (e.g., genes or transcripts). The GraphBG framework for unimodal spatial transcriptomics integrates graph-based learning with Bayesian statistics to identify spatial domains. The pipeline consists of the following key steps:

#### Data preprocessing

Similar to GraphST, GraphBG first processes the raw gene expression counts by performing library size normalization and log transformation using the Scanpy package [48]. The normalized gene expression matrix is then scaled to have zero mean and unit variance. Subsequently, the top 3000 highly variable genes (HVGs) are selected for downstream analysis. For datasets containing fewer than 3000 genes, the HVG selection step is skipped. Next, GraphBG applies Principal Component Analysis (PCA) to the normalized gene expression matrix to extract a compact, low-dimensional representation of the data. This transformation yields the PCA embedding *X*_emb_ ∈ ℝ^*N ×D*^, where *D* is the number of principal components. PCA projects the high-dimensional expression profiles onto a set of orthogonal directions that capture the greatest variance in the data. This step serves to denoise the input and reduce computational complexity in downstream steps.

#### Construction of neighbor graph using spatial location

To incorporate spatial context into the analysis, GraphBG constructs an undirected spatial neighbor graph *G* = (*V, E*), where *V* represents the set of *N* spatial spots and *E* encodes neighborhood relationships based on physical proximity. Each node in the graph corresponds to a spot, and edges connect spots that are spatially adjacent.

We define the adjacency matrix *A* ∈ ℝ^*N ×N*^ such that:

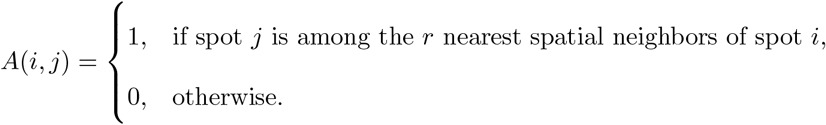

Here, spatial distance is measured using Euclidean distance between spot coordinates. The choice of *r* controls the local neighborhood size and determines the graph’s connectivity structure. In all experiments, we set *r* = 4, ensuring that each spot is connected to its four closest neighbors. This fixed neighborhood size provides a balance between capturing sufficient spatial context and avoiding over-smoothing or introducing long-range connections that may obscure local biological structures. The resulting graph *G* serves as the backbone for spatially-aware computations, including spectral graph convolutions in the next step.

#### First-order Chebyshev Approximation of Spectral Graph Convolutions

Spectral graph convolution defines the convolution of a graph signal *x* ∈ ℝ^*N*^ with a filter *g*_*θ*_ as:

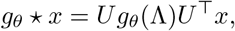

where *U* ∈ ℝ^*N ×N*^ is the matrix of eigenvectors of the normalized graph Laplacian *L* = *I* − *D*^−1*/*2^*AD*^−1*/*2^, Λ is the diagonal matrix of eigenvalues of *L*, and *g*_*θ*_(Λ) is a spectral filter function applied to the eigenvalues. Direct computation is computationally prohibitive for large graphs. To address this, the filter *g*_*θ*_(Λ) can be approximated using Chebyshev polynomials *T*_*k*_(*x*) up to order *K* [49]:

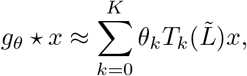

where 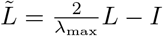 is the rescaled Laplacian so that its spectrum lies in [−1, 1], and *θ*_*k*_ are Chebyshev coefficients.

For the first-order approximation (*K* = 1), and assuming *λ*_max_ ≈ 2, the above simplifies to:

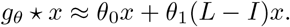

In particular, by choosing *θ*_0_ = 0, *θ*_1_ = −1, we obtain a simplified linear filter:

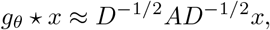

which corresponds to applying one step of a normalized graph smoothing operator to the signal *x*. Applying the above formulation to the PCA spot embeddings *X*_emb_, the graph convolution operation used in GraphBG is given by:

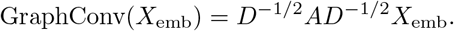

#### Variational Inference for Gaussian Mixture Models

Gaussian Mixture Models (GMMs) are probabilistic models that assume the data are generated from a mixture of Gaussian distributions, each associated with a latent cluster. Let *X* := GraphConv(*X*_emb_) = *{x*_1_, …, *x*_*N*_ *}* ⊂ ℝ^*D*^ denote the observed data obtained from the previous spectral graph convolutions, and *Z* = *{z*_1_, …, *z*_*N*_ *}* the set of latent variables, where each *z*_*n*_ is a 1-of-*K* binary indicator vector denoting cluster membership, and *K* is the number of clusters.

The likelihood function of the standard GMM is given by:

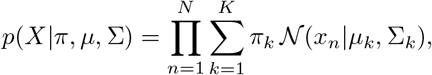

where *π*_*k*_ are the mixing coefficients, *µ*_*k*_ the mean vectors, and Σ_*k*_ the covariance matrices for each Gaussian component.

#### Maximum Likelihood Estimation and EM Algorithm

The model parameters are typically estimated by maximizing the log-likelihood:

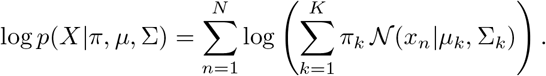

Since direct maximization is intractable, the Expectation-Maximization (EM) algorithm is commonly used. In the E-step, the posterior responsibilities *γ*(*z*_*nk*_) are computed, and in the M-step, the parameters (*π*_*k*_, *µ*_*k*_, Σ_*k*_) are re-estimated.

#### Bayesian Formulation

In the Bayesian GMM [23], priors are introduced on the model parameters. The mixing proportions follow a Dirichlet prior:

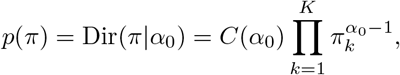

where *C*(*α*_0_) is the normalization constant.

For each component, the mean and precision are jointly governed by a Gaussian-Wishart prior:

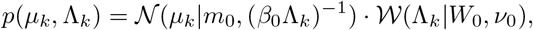

where 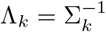 is the precision matrix, and (*m*_0_, *β*_0_, *W*_0_, *ν*_0_) are hyperparameters.

The joint distribution over all variables is:

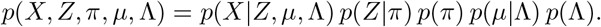

#### Variational Inference

Since the posterior *p*(*Z, π, µ*, Λ|*X*) is not analytically tractable, variational inference is used to approximate it with a factorized distribution:

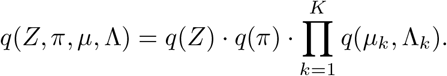

The variational distributions are optimized by maximizing the evidence lower bound (ELBO). The optimal form of *q*(*Z*) is:

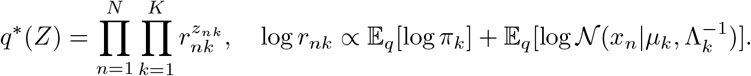

#### Variational EM Algorithm

The variational EM algorithm alternates between E-step and M-step [23]:

#### Variational E-step

Update responsibilities:

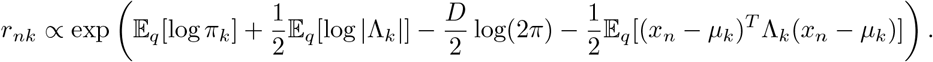

#### Variational M-step

Update variational distributions using sufficient statistics:

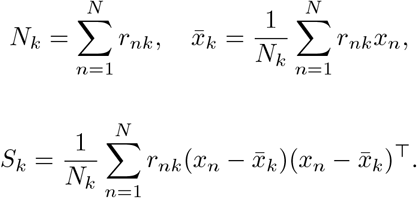

Updated Variational Distributions. Mixing proportions are given by

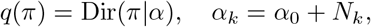

and means and precisions are given by

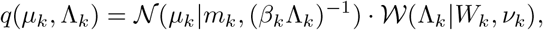

with updates:

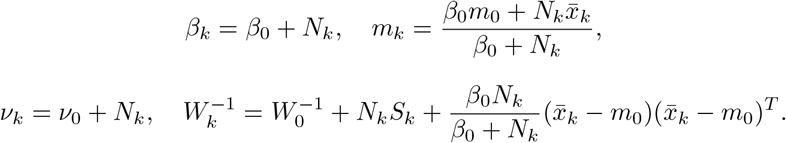

### Clustering Post-Processing

Each cluster inferred by the variational Bayesian Gaussian mixture model is interpreted as a spatial domain, consisting of spots with similar gene expression profiles and spatial proximity. Although the clustering integrates both expression and spatial information, misassignments can still occur, with some spots incorrectly assigned to domains that are spatially distant. To address this, GraphBG includes a post-processing refinement step. Specifically, for each spot *i*, the domain label is reassigned to match the most frequent label among its 50 nearest neighbors in spatial location space.

### 4.2. GraphBG-MS: Graph-Based Bayesian clustering for multi-slice Spatial Transcriptomics

While the previous section focused on identifying spatial domains within a single tissue slice, many spatial transcriptomics datasets span multiple slices or samples. These datasets pose additional challenges, including inter-slice variability, technical batch effects, and the need to align spatial domains across heterogeneous tissue sections. To address these complexities, we introduce *GraphBG-MS*, an extension of the single-slice GraphBG framework designed specifically for multi-slice spatial transcriptomics analysis.

GraphBG-MS integrates spatially aware representation learning with batch correction and Bayesian mixture modeling to enable robust and scalable clustering across multiple tissue slices. It is designed to preserve both transcriptomic similarity and spatial contiguity, while correcting for non-biological variations across slices. The GraphBG-MS pipeline consists of the following key stages:

#### Preprocessing

Raw gene expression matrices from each slice are normalized and log-transformed to stabilize variance and reduce the impact of sequencing depth and technical artifacts.

#### Graph Convolutional Encoding

Each tissue slice is treated as a spatial graph, where spots are nodes and edges encode spatial proximity. To embed local spatial context into expression features, we apply approximate spectral graph convolutions (as in GraphBG) to each slice independently, yielding spatially smoothed spot embeddings.

#### Metacell Construction

To reduce computational burden and enhance cross-slice comparability, each slice is aggregated into a fixed number of metacells (default *k* = 50) using MiniBatch k-Means clustering on the graph-convolved embeddings. Each metacell is represented by the average expression profile of its constituent spots.

#### Batch Effect Correction

Since slices may differ due to technical variation, we apply the ComBat algorithm [24] to the metacell expression profiles to remove batch effects while preserving biological structure. This harmonization step enables integration across slices for global clustering.

#### Global Clustering via Variational Inference

The batch-corrected metacell embeddings from all slices are jointly clustered using the variational Bayesian Gaussian mixture model [23].

#### Label Projection

Cluster labels assigned to metacells are projected back to their constituent spots, producing initial domain labels for all spots in all slices.

#### Spatial Refinement

To enhance spatial continuity and correct local misassignments, a refinement step is applied. For each spot *i*, its domain label is updated to match the most frequent label among its 50 nearest spatial neighbors. This local smoothing step encourages spatial coherence and improves domain boundary sharpness.

By combining spatial encoding, metacell abstraction, cross-slice correction, and probabilistic clustering, GraphBG-MS achieves accurate and coherent spatial domain identification across large-scale, multi-slice spatial transcriptomics datasets. The method provides a flexible framework that balances computational efficiency, statistical rigor, and spatial biological interpretability.

### 4.3. GraphBG-MM: multi-modal Spatial Transcriptomics Analysis

To extend our framework to multi-modal spatial transcriptomics (ST) data, we propose *GraphBG-MM*, which integrates spatial and cell molecular information from multiple modalities (e.g., gene expression and protein abundance) into a unified clustering framework. The pipeline proceeds through the following steps:

#### Preprocessing

Each modality is independently log-transformed and normalized, following the same preprocessing procedure as in the unimodal setting.

#### Graph Convolutional Encoding

For each modality, approximate spectral graph convolutions are applied independently to encode spatial context and produce modality-specific spatial embeddings.

#### multi-modal Integration via Kernel Multi-view CCA

To integrate spatial embeddings from multiple data modalities (e.g., gene expression, protein abundance), we employ *Kernel Multi-view Canonical Correlation Analysis (KCCA)* [25, 26]. Let 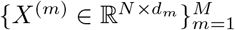 denote the set of spatial embeddings from *M* modalities, where each matrix *X*^(*m*)^ contains *N* shared samples (spots) and *d*_*m*_ features for modality *m*.

Each *X*^(*m*)^ is first mapped into a high-dimensional reproducing kernel Hilbert space (RKHS) using a positive-definite kernel *k*^(*m*)^(·, ·), such as the radial basis function (RBF) kernel. This results in a kernel matrix *K*^(*m*)^ ∈ ℝ^*N ×N*^, where 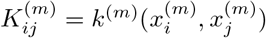, which is then centered.

KCCA seeks a set of projection directions *α*^(1)^, …, *α*^(*M*)^, one for each modality, that solve the following optimization problem:

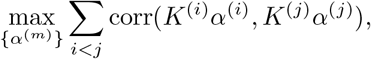

subject to the normalization constraints:

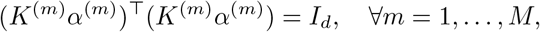

where *α*^(*m*)^ ∈ ℝ^*N ×d*^, and *d* is the number of canonical components retained.

After solving the optimization problem, each modality is projected into the shared latent space:

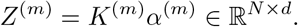

The final joint embedding *Z* ∈ ℝ^*N ×*(*M* ·*d*)^ is obtained by *concatenating* the projected views:

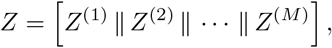

where ∥ denotes horizontal concatenation. This integrated representation captures modality-specific structure while aligning them in a common canonical space, and is used as input for downstream clustering.

#### Bayesian Clustering

The joint embedding is then clustered using a Variational Bayesian Gaussian Mixture model, enabling assignment of spots to spatial domains.

#### Spatial Refinement

A spatial smoothing step is applied, where each spot is reassigned to the most frequent domain label among its 50 nearest spatial neighbors, as in the unimodal GraphBG refinement step.

## Data availability

Unimodal and multi-slice ST datasets were obtained from Yuan, Z., Zhao, F., Lin, S. et al. Benchmarking spatial clustering methods with spatially resolved transcriptomics data. Nat Methods 21, 712–722 (2024). (https://figshare.com/projects/SDMBench/163942).

Multi-modal ST datasets were obtained from Long, Y., Ang, K.S., Sethi, R. et al. Deciphering spatial domains from spatial multi-omics with SpatialGlue. Nat Methods 21, 1658–1667 (2024). (https://github.com/JinmiaoChenLab/SpatialGlue). Mouse liver datasets were obtained from Cho et al. Microscopic examination of spatial transcriptome using Seq-Scope. Cell 184, 3559-3572.e22 (2021).

## Code availability

The GraphBG software is available at GitHub (https://github.com/CamiLQDTULab/GraphBG) under the open-source BSD license. The GraphBG repository also includes all code necessary to reproduce the results of this manuscript.

## Acknowledgements

V.H. Do gratefully acknowledges the hospitality and financial support provided by Vietnam Institute for Advanced Studies in Mathematics (VIASM) through the *IAPB* program.

We would like to thank Dongyuan Song for his generous help with the scDesign3 simulation, which significantly facilitated our analysis.

We thank the authors of *scNiche* for providing the Python implementation, which facilitated our analysis of cellular niches.

## Author’s Contributions

V.H.D. and S.C. conceptualize and designed the project. V.H.D. and T.P.L.T. performed data analysis and interpretation. V.H.D. developed and implemented GraphBG and prepared documentation. All authors wrote the paper and commented on the manuscript at all stages.

## References

[1] Vandereyken, K., Sifrim, A., Thienpont, B., Voet, T.: Methods and applications for single-cell and spatial multi-omics. Nature Reviews Genetics 24(8), 494–515 (2023). doi:10.1038/s41576-023-00580-2

[2] Seferbekova, Z., Lomakin, A., Yates, L.R., Gerstung, M.: Spatial biology of cancer evolution. Nature Reviews Genetics 24(5), 295–313 (2022). doi:10.1038/s41576-022-00553-x

[3] Chen, W.-T., Lu, A., Craessaerts, K., Pavie, B., Sala Frigerio, C., Corthout, N., Qian, X., Laláková, J., Kühnemund, M., Voytyuk, I., Wolfs, L., Mancuso, R., Salta, E., Balusu, S., Snellinx, A., Munck, S., Jurek, A., Fernandez Navarro, J., Saido, T.C., Huitinga, I., Lundeberg, J., Fiers, M., De Strooper, B.: Spatial transcriptomics and in situ sequencing to study alzheimer’s disease. Cell 182(4), 976–99119 (2020). doi:10.1016/j.cell.2020.06.038

[4] Stahl, P.L., Salmen, F., Vickovic, S., et al., Frisén, J.: Visualization and analysis of gene expression in tissue sections by spatial transcriptomics. Science 353(6294), 78–82 (2016). doi:10.1126/science.aaf2403

[5] Yang, W., Wang, P., Xu, S., Wang, T., Luo, M., Cai, Y., Xu, C., Xue, G., Que, J., Ding, Q., Jin, X., Yang, Y., Pang, F., Pang, B., Lin, Y., Nie, H., Xu, Z., Ji, Y., Jiang, Q.: Deciphering cell–cell communication at single-cell resolution for spatial transcriptomics with subgraph-based graph attention network. Nature Communications 15(1) (2024). doi:10.1038/s41467-024-51329-2

[6] Shi, H., He, Y., Zhou, Y., Huang, J., Maher, K., Wang, B., Tang, Z., Luo, S., Tan, P., Wu, M., Lin, Z., Ren, J., Thapa, Y., Tang, X., Chan, K.Y., Deverman, B.E., Shen, H., Liu, A., Liu, J., Wang, X.: Spatial atlas of the mouse central nervous system at molecular resolution. Nature 622(7983), 552–561 (2023). doi:10.1038/s41586-023-06569-5

[7] Zhang, M., Eichhorn, S.W., Zingg, B., Yao, Z., Cotter, K., Zeng, H., Dong, H., Zhuang, X.: Spatially resolved cell atlas of the mouse primary motor cortex by merfish. Nature 598(7879), 137–143 (2021). doi:10.1038/s41586-021-03705-x

[8] Zhang, M., Pan, X., Jung, W., Halpern, A.R., Eichhorn, S.W., Lei, Z., Cohen, L., Smith, K.A., Tasic, B., Yao, Z., Zeng, H., Zhuang, X.: Molecularly defined and spatially resolved cell atlas of the whole mouse brain. Nature 624(7991), 343–354 (2023). doi:10.1038/s41586-023-06808-9

[9] Yuan, Z., Zhao, F., Lin, S., Zhao, Y., Yao, J., Cui, Y., Zhang, X.-Y., Zhao, Y.: Benchmarking spatial clustering methods with spatially resolved transcriptomics data. Nature Methods 21(4), 712–722 (2024). doi:10.1038/s41592-024-02215-8

[10] Hu, Y., Xie, M., Li, Y., Rao, M., Shen, W., Luo, C., Qin, H., Baek, J., Zhou, X.M.: Benchmarking clustering, alignment, and integration methods for spatial transcriptomics. Genome Biology 25(1) (2024). doi:10.1186/s13059-024-03361-0

[11] Dries, R., Chen, J., del Rossi, N., Khan, M.M., Sistig, A., Yuan, G.-C.: Advances in spatial transcriptomic data analysis. Genome Research 31(10), 1706–1718 (2021). doi:10.1101/gr.275224.121

[12] Pham, D., Tan, X., Balderson, B., Xu, J., Grice, L.F., Yoon, S., Willis, E.F., Tran, M., Lam, P.Y., Raghubar, A., Kalita-de Croft, P., Lakhani, S., Vukovic, J., Ruitenberg, M.J., Nguyen, Q.H.: Robust mapping of spatiotemporal trajectories and cell–cell interactions in healthy and diseased tissues. Nature Communications 14(1) (2023). doi:10.1038/s41467-023-43120-6

[13] Dong, K., Zhang, S.: Deciphering spatial domains from spatially resolved transcriptomics with an adaptive graph attention auto-encoder. Nature Communications 13(1) (2022). doi:10.1038/s41467-022-29439-6

[14] Blondel, V.D., Guillaume, J.-L., Lambiotte, R., Lefebvre, E.: Fast unfolding of communities in large networks. Journal of Statistical Mechanics: Theory and Experiment 2008(10), 10008 (2008). doi:10.1088/1742-5468/2008/10/P10008

[15] Satija, R., Farrell, J.A., Gennert, D., Schier, A.F., Regev, A.: Spatial reconstruction of single-cell gene expression data. Nature Biotechnology 33(5), 495–502 (2015). doi:10.1038/nbt.3192

[16] Hu, J., Li, X., Coleman, K., Schroeder, A., Ma, N., Irwin, D.J., Lee, E.B., Shinohara, R.T., Li, M.: Spagcn: Integrating gene expression, spatial location and histology to identify spatial domains and spatially variable genes by graph convolutional network. Nature Methods 18(11), 1342–1351 (2021). doi:10.1038/s41592-021-01255-8

[17] Zhao, E., Stone, M.R., Ren, X., Guenthoer, J., Smythe, K.S., Pulliam, T., Williams, S.R., Uytingco, C.R., Taylor, S.E.B., Nghiem, P., Bielas, J.H., Gottardo, R.: Spatial transcriptomics at subspot resolution with bayesspace. Nature Biotechnology 39(11), 1375–1384 (2021). doi:10.1038/s41587-021-00935-2

[18] Li, J., Chen, S., Pan, X., Yuan, Y., Shen, H.-B.: Cell clustering for spatial transcriptomics data with graph neural networks. Nature Computational Science 2(6), 399–408 (2022). doi:10.1038/s43588-022-00266-5

[19] Ren, H., Walker, B.L., Cang, Z., Nie, Q.: Identifying multicellular spatiotemporal organization of cells with spaceflow. Nature Communications 13(1) (2022). doi:10.1038/s41467-022-31739-w

[20] Long, Y., Ang, K.S., Li, M., Chong, K.L.K., Sethi, R., Zhong, C., Xu, H., Ong, Z., Sachaphibulkij, K., Chen, A., Zeng, L., Fu, H., Wu, M., Lim, L.H.K., Liu, L., Chen, J.: Spatially informed clustering, integration, and deconvolution of spatial transcriptomics with graphst. Nature Communications 14(1) (2023). doi:10.1038/s41467-023-36796-3

[21] Korsunsky, I., Millard, N., Fan, J., Slowikowski, K., Zhang, F., Wei, K., Baglaenko, Y., Brenner, M., Loh, P.-r., Raychaudhuri, S.: Fast, sensitive and accurate integration of single-cell data with harmony. Nature Methods 16(12), 1289–1296 (2019). doi:10.1038/s41592-019-0619-0

[22] Long, Y., Ang, K.S., Sethi, R., Liao, S., Heng, Y., van Olst, L., Ye, S., Zhong, C., Xu, H., Zhang, D., Kwok, I., Husna, N., Jian, M., Ng, L.G., Chen, A., Gascoigne, N.R.J., Gate, D., Fan, R., Xu, X., Chen, J.: Deciphering spatial domains from spatial multi-omics with spatialglue. Nature Methods 21(9), 1658–1667 (2024). doi:10.1038/s41592-024-02316-4

[23] Bishop, C.M.: Pattern Recognition and Machine Learning. Springer, New York (2006)

[24] Zhang, Y., Parmigiani, G., Johnson, W.E.: Combat-seq: batch effect adjustment for rna-seq count data. NAR Genomics and Bioinformatics 2(3) (2020). doi:10.1093/nargab/lqaa078

[25] Hardoon, D.R., Szedmak, S., Shawe-Taylor, J.: Canonical correlation analysis: An overview with application to learning methods. Neural Computation 16(12), 2639–2664 (2004). doi:10.1162/0899766042321814

[26] Bach, F.R., Jordan, M.I.: Kernel independent component analysis. Journal of Machine Learning Research 3, 1–48 (2002)

[27] Li, Z., Zhou, X.: Bass: multi-scale and multi-sample analysis enables accurate cell type clustering and spatial domain detection in spatial transcriptomic studies. Genome Biology 23(1) (2022). doi:10.1186/s13059-022-02734-7

[28] Cang, Z., Ning, X., Nie, A., Xu, M., Zhang, J.: Scan-it: Domain segmentation of spatial transcriptomics images by graph neural network. Proceedings of the British Machine Vision Conference (BMVC) (406) (2021)

[29] Maynard, K.R., Collado-Torres, L., Weber, L.M., Uytingco, C., Barry, B.K., Williams, S.R., Catallini, J.L., Tran, M.N., Besich, Z., Tippani, M., Chew, J., Yin, Y., Kleinman, J.E., Hyde, T.M., Rao, N., Hicks, S.C., Martinowich, K., Jaffe, A.E.: Transcriptome-scale spatial gene expression in the human dorsolateral prefrontal cortex. Nature Neuroscience 24(3), 425–436 (2021). doi:10.1038/s41593-020-00787-0

[30] Chen, X., Sun, Y.-C., Church, G.M., Lee, J.H., Zador, A.M.: Efficient in situ barcode sequencing using padlock probe-based baristaseq. Nucleic Acids Research 46(4), 22–22 (2017). doi:10.1093/nar/gkx1206

[31] Chen, K.H., Boettiger, A.N., Moffitt, J.R., Wang, S., Zhuang, X.: Spatially resolved, highly multiplexed rna profiling in single cells. Science 348(6233) (2015). doi:10.1126/science.aaa6090

[32] Codeluppi, S., Borm, L.E., Zeisel, A., La Manno, G., van Lunteren, J.A., Svensson, C.I., Linnarsson, S.: Spatial organization of the somatosensory cortex revealed by osmfish. Nature Methods 15(11), 932–935 (2018). doi:10.1038/s41592-018-0175-z

[33] Wang, X., Allen, W.E., Wright, M.A., Sylwestrak, E.L., Samusik, N., Vesuna, S., Evans, K., Liu, C., Ramakrishnan, C., Liu, J., Nolan, G.P., Bava, F.-A., Deisseroth, K.: Three-dimensional intact-tissue sequencing of single-cell transcriptional states. Science 361(6400) (2018). doi:10.1126/science.aat5691

[34] Song, D., Wang, Q., Yan, G., Liu, T., Sun, T., Li, J.J.: scdesign3 generates realistic in silico data for multimodal single-cell and spatial omics. Nature Biotechnology 42(2), 247–252 (2023). doi:10.1038/s41587-023-01772-1

[35] Stoeckius, M., Hafemeister, C., Stephenson, W., Houck-Loomis, B., Chattopadhyay, P.K., Swerdlow, H., Satija, R., Smibert, P.: Simultaneous epitope and transcriptome measurement in single cells. Nature Methods 14(9), 865–868 (2017). doi:10.1038/nmeth.4380

[36] Desikan, R.S., Ségonne, F., Fischl, B., Quinn, B.T., Dickerson, B.C., Blacker, D., Buckner, R.L., Dale, A.M., Maguire, R.P., Hyman, B.T., Albert, M.S., Killiany, R.J.: An automated labeling system for subdividing the human cerebral cortex on mri scans into gyral based regions of interest. NeuroImage 31(3), 968–980 (2006). doi:10.1016/j.neuroimage.2006.01.021

[37] Liao, S., Heng, Y., Liu, W., Xiang, J., Ma, Y., Chen, L., Feng, X., Jia, D., Liang, D., Huang, C., Zhang, J., Jian, M., Su, K., Li, M., Loh, Y.H., Chen, A., Xu, X.: Integrated spatial transcriptomic and proteomic analysis of fresh frozen tissue based on stereo-seq (2023). doi:10.1101/2023.04.28.538364

[38] Ben-Chetrit, N., Niu, X., Swett, A.D., Sotelo, J., Jiao, M.S., Stewart, C.M., Potenski, C., Mielinis, P., Roelli, P., Stoeckius, M., Landau, D.A.: Integration of whole transcriptome spatial profiling with protein markers. Nature Biotechnology 41(6), 788–793 (2023). doi:10.1038/s41587-022-01536-3

[39] He, S., Bieniosek, M., Song, D., Zhou, J., Chidester, B., Wu, Z., Boen, J., Sharma, P., Trevino, A.E., Zou, J.: Learning single-cell spatial context through integrated spatial multiomics with coral. bioXiv (2025). doi:10.1101/2025.02.01.636038

[40] Qian, J., Shao, X., Bao, H., Fang, Y., Guo, W., Li, C., Li, A., Hua, H., Fan, X.: Identification and characterization of cell niches in tissue from spatial omics data at single-cell resolution. Nature Communications 16(1) (2025). doi:10.1038/s41467-025-57029-9

[41] Allen, W.E., Blosser, T.R., Sullivan, Z.A., Dulac, C., Zhuang, X.: Molecular and spatial signatures of mouse brain aging at single-cell resolution. Cell 186(1), 194–20818 (2023). doi:10.1016/j.cell.2022.12.010

[42] Cho, C.-S., Xi, J., Si, Y., Park, S.-R., Hsu, J.-E., Kim, M., Jun, G., Kang, H.M., Lee, J.H.: Microscopic examination of spatial transcriptome using seq-scope. Cell 184(13), 3559–357222 (2021). doi:10.1016/j.cell.2021.05.010

[43] Halpern, K.B., Shenhav, R., Matcovitch-Natan, O., Tóth, B., Lemze, D., Golan, M., Massasa, E.E., Baydatch, S., Landen, S., Moor, A.E., Brandis, A., Giladi, A., Stokar-Avihail, A., David, E., Amit, I., Itzkovitz, S.: Single-cell spatial reconstruction reveals global division of labour in the mammalian liver. Nature 542(7641), 352–356 (2017). doi:10.1038/nature21065

[44] Ben-Moshe, S., Shapira, Y., Moor, A.E., Manco, R., Veg, T., Bahar Halpern, K., Itzkovitz, S.: Spatial sorting enables comprehensive characterization of liver zonation. Nature Metabolism 1(9), 899–911 (2019). doi:10.1038/s42255-019-0109-9

[45] Richter, M.L., Deligiannis, I.K., Yin, K., Danese, A., Lleshi, E., Coupland, P., Vallejos, C.A., Matchett, K.P., Henderson, N.C., Colome-Tatche, M., Martinez-Jimenez, C.P.: Single-nucleus rna-seq2 reveals functional crosstalk between liver zonation and ploidy. Nature Communications 12(1) (2021). doi:10.1038/s41467-021-24543-5

[46] Hildebrandt, F., Andersson, A., Saarenpää, S., Larsson, L., Van Hul, N., Kanatani, S., Masek, J., Ellis, E., Barragan, A., Mollbrink, A., Andersson, E.R., Lundeberg, J., Ankarklev, J.: Spatial transcriptomics to define transcriptional patterns of zonation and structural components in the mouse liver. Nature Communications 12(1) (2021). doi:10.1038/s41467-021-27354-w

[47] Wang, S., Wang, X., Shan, Y., Tan, Z., Su, Y., Cao, Y., Wang, S., Dong, J., Gu, J., Wang, Y.: Region-specific cellular and molecular basis of liver regeneration after acute pericentral injury. Cell Stem Cell 31(3), 341–3587 (2024). doi:10.1016/j.stem.2024.01.013

[48] Wolf, F.A., Angerer, P., Theis, F.J.: Scanpy: large-scale single-cell gene expression data analysis. Genome Biology 19(1) (2018). doi:10.1186/s13059-017-1382-0

[49] Defferrard, M., Bresson, X., Vandergheynst, P.: Convolutional neural networks on graphs with fast localized spectral filtering. In: Proceedings of the 30th International Conference on Neural Information Processing Systems. NIPS’16, pp. 3844–3852. Curran Associates Inc., Red Hook, NY, USA (2016)

